# Higher-Order Interaction Analysis via Hypergraph Models for Studying Multidimensional Neuroscience Data

**DOI:** 10.1101/2024.11.22.624800

**Authors:** Dalma Bilbao, Hugo Aimar, Pablo Torterolo, Diego M. Mateos

## Abstract

Higher-Order Interaction (HOI) theory offers a powerful framework for capturing complex, non-linear relationships within multidimensional systems, moving beyond traditional pairwise graph methods to encompass multi-way interactions. This study applies HOI analysis, specifically using hypergraph theory, to explore intricate connectivity patterns in electrophysiological signals from neuroscience. Hypergraphs were constructed from connectivity data across various frequency bands, characterized through metrics such as spectral entropy, hyperedge centrality, and vertex centrality, and compared using spectral and centrality distance measures. Three distinct neurophysiological datasets were analyzed: intracranial EEG signals from rats during different sleep stages, scalp EEG data to distinguish between epilepsy types, and MEG recordings of seizure dynamics. The findings highlight the effectiveness of hypergraph-based HOI analysis in mapping neural dynamics across normal and pathological brain states. In sleep studies, it reveals distinct connectivity patterns between REM and NREM stages, while in epilepsy, it differentiates seizure types and stages, identifying spectral entropy as a potential marker for seizure onset. Notably, HOI analysis captures differences between primary and secondary generalized epilepsy, suggesting enhanced diagnostic accuracy. This approach provides a powerful tool for understanding complex neural interactions in high-dimensional data.

## 1 Introduction

In order to study the interrelationship between components of a system, the most commonly used analyses are derived from graph theory, which is based on pairwise relationships between components. However, in the vast majority of natural systems, the relationships between components extend beyond simple pairs, manifesting in a more intricate manner. This underscores the necessity for alternative approaches to elucidate the system’s underlying complexities. One such theory that attempted to address this challenge is that of Higher-Order Interactions (HOI), which seeks to capture the intricate, non-linear interactions within systems.

Higher-order interaction analysis offers several key advantages for understanding complex systems. Unlike traditional methods, it captures multi-way interactions that go beyond simple pairwise relationships. This broader scope improves predictive accuracy and reveals hidden, non-linear patterns in the data. As a result, HOI is particularly useful in fields such as machine learning, neuroscience, genomics and social network analysis, where the complexity of the data often requires more refined models[1]. By addressing high-dimensional data and modeling intricate dependencies, high-order analysis improves model robustness and flexibility while offering more profound and nuanced insights that are often overlooked by traditional methods.

While Higher-Order Interaction (HOI) theory has shown promise across various fields, its impact is particularly evident in neuroscience, where the complexity of brain activity demands sophisticated analytical tools. Advances in neuroscience technology have enabled the collection of vast and multidimensional datasets, such as high-density electroencephalograms (hEEG), magnetoencephalograms (MEG), and high-density multielectrode recordings (hMER). These technologies generate a substantial quantity of high-dimensional data. However, the exponential growth in the volume of data makes the process of analysis increasingly challenging. In this context, methods based on HOI theory emerge as highly effective tools for addressing this challenge. Currently, HOI analysis has been implemented in the study of neuroscientific signals. For example, it has been used to investigate major depressive disorder [2], neurodegenerative diseases such as Alzheimer’s [3], the interplay between aging and cognitive function [4], and in the diagnosis of autism spectrum disorders [5].

One of the most widely used approaches to studying higher-order interaction in various systems is through Hypergraph Theory. This provides a robust and flexible framework for understanding complex, multi-dimensional relationships across diverse domains [6]. In contrast to traditional graphs, where edges connect only two vertices, hypergraphs generalise this concept by allowing hyperedges to connect multiple vertices simultaneously. This enables a more natural and effective modelling of group interactions. This feature makes hypergraphs particularly useful for capturing interactions at higher levels of complexity, such as collective dynamics in biological [7, 8, 9], medical [10], social [11, 12], and technological systems [13, 14]. Particularly in the field of neuroscience, hypergraph theory has been employed to capture real interaction relationships among brain regions, improving the accuracy of diagnosing autism spectrum disorders [5], classifying and subtyping psychiatric disorders [15], and aiding in migraine classification [16].

In this study, we introduce a method for analyzing higher-order interactions in multidimensional electrophysiological signals using hypergraph theory. We construct hypergraphs based on simple graphs derived from connectivity analyses across various electrophysiological frequency bands, with connectivity measured through the Phase Locking Index (PLI) [17]. Each hypergraph is then characterized by three metrics rooted in hypergraph theory: (i) spectral entropy, (ii) hyperedge centrality, and (iii) vertex centrality. Additionally, we define three dissimilarity metrics to compare hypergraphs: (i) spectral distance, (ii) hyperedge centrality distance, and (iii) vertex centrality distance.

To evaluate our approach, we applied these hypergraph generation, quantification, and comparison methods to three distinct examples of multidimensional neurophysiological signals. First, we examined and characterized intracranial electroencephalogram (iEEG) signals from rats across different sleep stages. Second, we analyzed scalp electroencephalogram (EEG) signals to differentiate between various types of epilepsy in human patients. Finally, we investigated the temporal evolution of neuronal network dynamics in magnetoencephalography (MEG) recording from two patients during epileptic seizures.

Our multilayer hypergraph analysis method demonstrates the capability to capture higher-order interactions in multidimensional electrophysiological signals. This approach enables the study and comparison of underlying neural network dynamics across normal and pathological brain states.

## 2 Introduction Hypergraphs and Their Quantifiers

### 2.1 Hypergraphs

Hypergraphs are generalizations of graphs such that each edge is allowed to contain more than two vertices. Formally, an (undirected) hypergraph is a pair ℋ = (𝒱, ℰ) where 𝒱 is the set of vertices and ℰ is a subset of non-empty parts of 𝒱 that cover 𝒱. The elements of ℰ are called hyperedges [18]. In other words, *e*≠∅ for all *e* ∈ ℰ and ∪_*e* ∈ ℰ_ *e* = 𝒱

The **order** of the hypergraph ℋ = (𝒱, ℰ) is the number of the elements of 𝒱, denoted as |𝒱| = *n*, and its **size** is the number of element of ℰ, denoted as |ℰ| = *m*.

Given a hypergraph ℋ = (𝒱, ℰ), the **incidence matrix** *H* of order *n* × *m* associated with ℋ is defined as

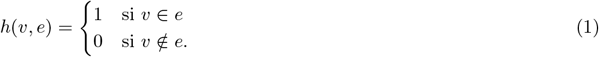

From the incidence matrix *H*, we can calculate two diagonal matrices to represent the **vertex degrees** as follows

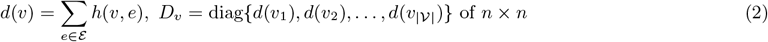

and **hyperedge degrees**

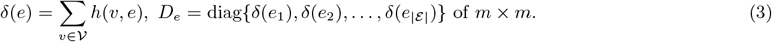

The **adjacency matrix** 𝒜(ℋ) of ℋ is defined as a square matrix *n* × *n*, such that 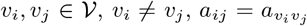 is the number of hyperedges containing both *v*_*i*_ and *v*_*j*_, in other words

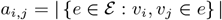

and *a*_*i,i*_ = 0. This matrix is symmetric, and all *a*_*ij*_ are non-negative integers.

So that the hypergraph ℋ induces a weighted undirected graph in the same set of vertices. Then we have a well-defined **Laplacian matrix** ℒ(ℋ), of size *n* × *n*, given by

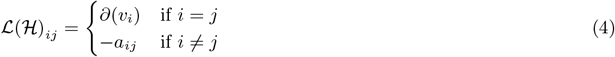

where 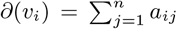 Since ℒ(ℋ) is symmetric and positive semidefinite, then ℒ(ℋ) as a spectral resolution with non-negative eigenvalues 0 = *µ*_0_ ≤ … ≤ *µ*_*n*_, the smallest eigenvalue is 0 = *µ*_0_. Moreover,

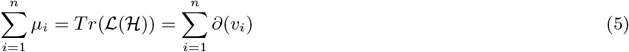

where *T r*(ℒ(ℋ)) is the trace of the matrix ℒ(ℋ). In matrix form ℒ(ℋ) = *∂* − 𝒜(ℋ), where *∂* = *diag*(*∂*(*v*_1_), …*∂*(*v*_*n*_)).

### 2.2. Hypergraph quatifiers

To effectively extract features from hypergraphs, it is essential to quantify their properties. This work focuses on three key quantifiers: entropy derived from the Laplacian spectrum, vertex centrality, and hyperedge centrality.

#### 2.2.1. Spectral Entropy

To introduce the analogue of the notion of Von Neumann entropy [19], we first define the following normalization of the Laplacian

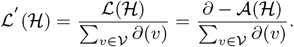

The eigenvalues of this matrix are

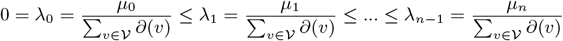

for all *i* ∈ {1, 2, …*n*}, while the eigenvectors are the same as those of ℒ.

The set of eigenvalues {*λ*_*i*_} with *i* ∈ {1, 2, …, *n*} satisfies:

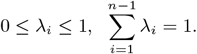

Because of that, this set can be seen as a discrete probability distribution. Hence, we can define the algebraic hypergraph entropy S(ℋ) by

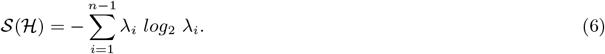

This notion of entropy is associated to a given hypergraph through the weighted undirected graph built through its adjacency matrix. As usual, the notion of entropy S(ℋ) is a measure of the information contained in the higher order interactions described by ℋ.

#### 2.2.2 Vertices and hyperedges centralities

In the case of simple graphs, centrality refers to a measure of how important a vertex (or node) is within the network represented by G = (𝒱, ℰ). The simplest way to define the centrality of a vertex in a simple graph is by counting the number of edges connected to that vertex. This concept can be easily extended to hypergraphs. Let ℋ = (𝒱, ℰ) be a given hypergraph. The function 𝒞_V_ : 𝒱 → ℕ defined by

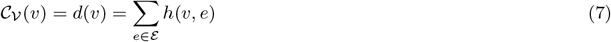

is called the **vertex degree centrality** of ℋ. This function captures the significance of each node in the hypergraph ℋ within the connectivity structure defined by ℋ.

**Hyperedge centrality** indicates how critical a particular hyperedge is to the overall connectivity of the hypergraph ℋ. In the similar way as before, we can define a function 𝒞_E_ : ℰ → ℕ defined by

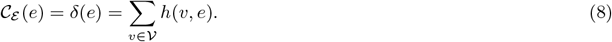

providing valuable information about the structure and dynamics of the relationships in a hipergraph.

### 2.3 Distances between hypergraphs

The literature on graph dissimilarity measures is extensive [20], with several approaches defining natural distances between hypergraphs by utilizing the weighted undirected graph derived from the adjacency matrix *A*(ℋ). This framework is particularly useful for defining spectral distances between hypergraphs that have the same number of vertices. In this work, we introduce three types of distance (or dissimilarity) measures for hypergraphs, based on the previously described quantifiers.

#### 2.3.1 Spectral distance

Given two hypergraphs ℋ = (𝒱, ℰ) and 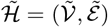 with 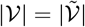, let 𝒜 and 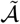denote their respective adjacency matrices, and ℒ and 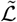 denote the associated normalized Laplacians. The spectral decompositions of ℒ and 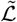 yield the eigenvalues 0 = *λ*_0_ ≤ *λ*_1_ ≤ … ≤ *λ*_*n*−1_ and 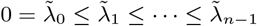, respectively. These two sequences, regarded as vectors in ℝ^*n*−1^, have defined *p*-distances, 1 ≤ *p <* ∞.

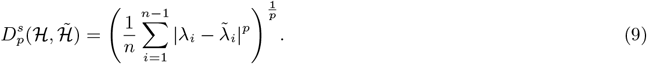

The most important case is *p* = 2 that well defines the Hilbert space structure in ℝ^*n*−1^.

#### 2.3.2. Vertex centrality distance

Given hypergraphs ℋ and, 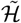 as before, now assume that 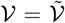. Let 𝒞_𝒱_ and 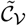 denote the respective vertex centrality functions. A dissimilarity measure between ℋ and 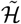 that accounts for vertex centrality is defined by

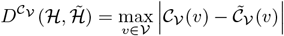

Notice that, from the definition of 𝒞_V_ and 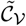, we have

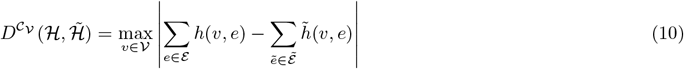

where *h* and 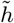 provide the respective incidence matrices for ℋ and 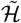.

#### 2.3.3 Hyperedge centrality distance

Similarly, if we have ℋ and 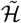 and assuming now that 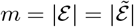 a hyperedge centrality-based distance between ℋ and 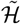 can be defined by

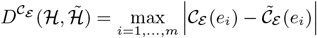

with the above notation, can be computed as

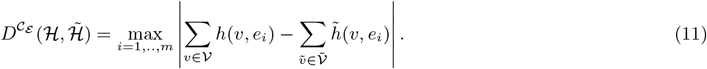

## 3 Hypergraph bulding

In this section, we introduce our method for constructing hypergraphs based on multilayer graphs. To achieve this, each layer or graph must share the same set of vertices *v* = {*v*_1_, …, *v*_*k*_}, however the set of edges *E* can vary depending on the relationships present between edges in each layer.

In particular, for this study, we use neurophysiological recording electrodes as edge, and the different layers or graphs are represented by the connectivity between those electrodes across different frequency bands B. It is important to note that this method is not restricted to this specific type of data, but can be generalized to generate hypergraphs for any problem that meets the conditions described above. For example, it could be applied to a map with fixed cities (nodes), where the different graphs or layers represent relationships between cities via different transportation methods, such as roads, railways, or flights.

For the case of neurophysiological signals such as electroencephalogram (EEG), magnetoencephalogram (MEG), or intracranial EEG (iEEG), the first step is to take the original signal (which may have been preprocessed) and filter it into the frequency bands relevant to the problem at hand. A general case would be to filter into the most neurophysiologically relevant bands, such as Theta, Delta, Alpha, Beta, and Gamma B = {*θ, δ, α, β, γ*}. Once filtered, the correlation between all pairs of signals is calculated. In our case, we applied a phase-based correlation measure known as the *Phase Locking Index* (PLI) [17], resulting in a *N*_*chan*_ × *N*_*chan*_ matrix 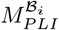 for each frequency band.

Since PLI values are continuous between 0 and 1, we binarize the matrix using a threshold matrix 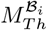. This step removes spurious connections. The threshold matrix 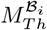 is calculated by first generating *N*_*sur*_ = 30 surrogate datasets from the filtered signals and then computing the PLI for each surrogate, obtaining 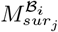 or *j* = 1, …, *N*_*sur*_. The threshold matrix is then calculated as 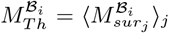 Finally, the binarized matrix 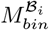 is obtained by comparing the original PLI matrix 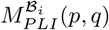 with the threshold matrix: if 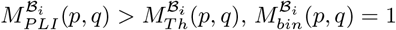, otherwise it is set to 0. One important point to consider when applying any metric that involves the Hilbert transform, such as PLI, is that the signal should be filtered in a narrow frequency band to extract the phase of the wave as cleanly as possible [21]. In our case, we used a frequency bandwidth of 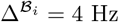.

Once the graphs or layers are obtained for each frequency band, we proceed to build the incidence matrix *h* of size *N*_*band*_ × *N*_*pair*_, where *N*_*band*_ is the number of frequency bands and *N*_*pair*_ is the number of possible connections between electrode pairs, 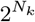. Each row of the incidence matrix corresponds to a frequency band, and entries are set to 0 or 1 depending on whether a given pair of electrodes is connected in the corresponding connectivity matrix. Once the incidence matrix *h* is constructed, we have the hypergraph associated with the multilayer graph and the adjacency matrix associa, and we can apply the quantifiers and distances introduced in section 2.2.Figure 1 provides a graphical overview of the pipeline used to derive hypergraphs from multilayer graphs.

**Figure 1:**
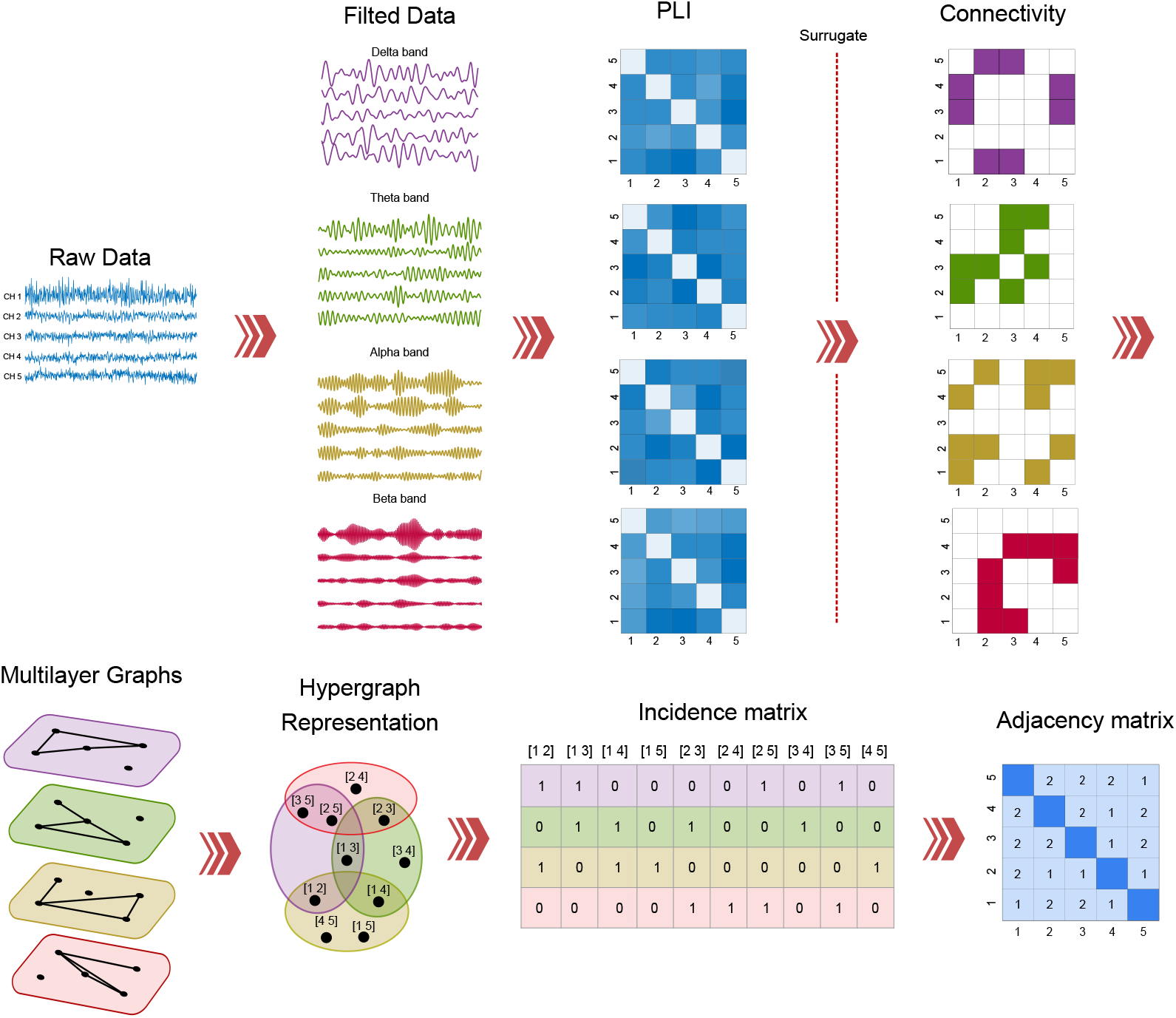
Pipeline for obtaining a hypergraph representation of high-order interactions from multilayer graphs in electrophysiological recordings

## 4 Results

The model of building hypergraphs and their quantifiers, as described above, was applied to three different examples. The first example is an analysis of intracranial EEG (iEEG) recordings in rats during different sleep states: Active Wakefulness (AW), Quiet Wakefulness (QW), and Non-Rapid Eye Movement (NREM), Rapid Eye Movement (REM). The second example presents a study of scalp EEG recordings from epileptic patients suffering from different types of epilepsy. Finally, the third example examines the dynamical networks changes in MEG signals from two patients with generalized epilepsy during seizures.

### 4.1 Quantifying sleep states

The initial example was focused on the study of brain dynamics through the analysis of higher-order interactions within the brain networks of rats across different sleep stages. We examine iEEG signals from 9 rats across the four distinct sleep states. The data were obtained by the Sleep Neurobiology Laboratory group at the University of the Republic in Uruguay, led by Dr. Torterolo. Each recording have 7 electrodes positioned in the following brain regions: V2 (secondary visual area), S1 (primary somatosensory cortex), M1 (primary motor cortex), and OB (olfactory bulb) (See Figure 2A, left panel). The first step was to apply a bandpass filter with a passband between 0.5 and 70 Hz. Additionally, a notch filter was employed to eliminate 60 Hz power line noise. Subsequently, epochs devoid of artefacts and exhibiting non-transitory characteristics were meticulously selected for quantitative EEG analysis. Spike detection algorithms were used to extracted epochs corresponding to each sleep state. Active Wakefulness (AW), Quiet Wakefulness (QW), Non-REM (NREM) sleep, and REM sleep states were manually scored in 3-second epochs using Spike 2 software (Cambridge Electronic Design), based on the criteria described in [22]. Particularly, Wakefullnes state were categorized into active wakefulness (AW) and quiet wakefulness (QW) based on the average theta activity recorded across four posterior electrodes. Epochs with theta power exceeding the 70th percentile were designated as AW, while the remaining epochs were classified as QW as done in these previous works [23, 24].

**Figure 2:**
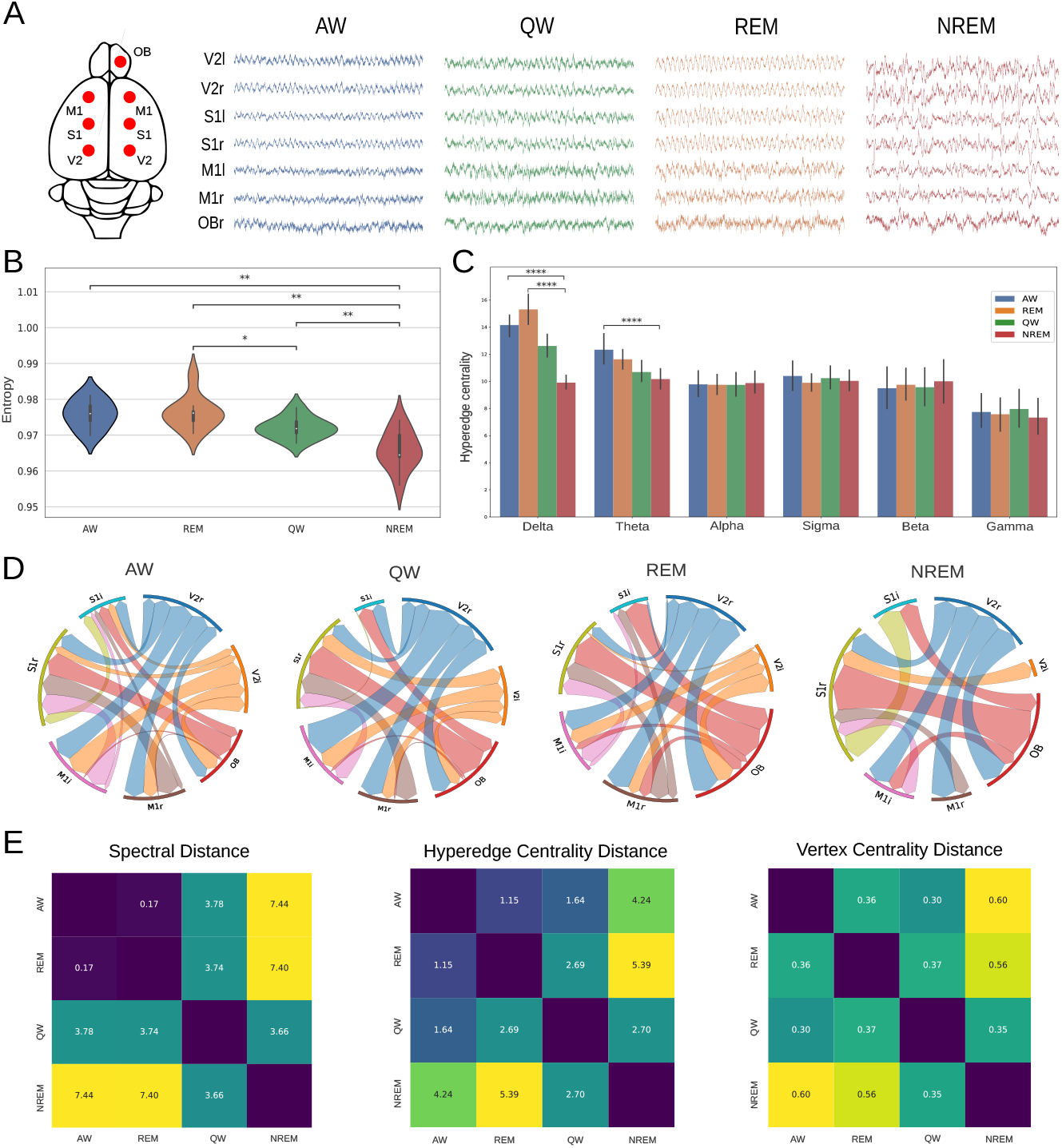
Study of Higher-Order Interactions (HOI) using hypergraph analysis in neural network dynamics across different sleep stages in rats. **A)** Topological recording positions in cortical areas in rats (left panel) and example raw data recordings across the four sleep stages (right panel). **B)** Spectral entropy analysis across different sleep states. The violin plot shows the average entropy values over 100 epochs for each rat and sleep state. Statistical comparisons between states were conducted using the Kruskal-Wallis test, with Dunn’s multiple comparisons test indicating significant differences: AW vs. NREM *∗ ∗ p* = 2×10*−*3, REM vs. QW *∗ p* = 3×10*−*2; REM vs. NREM *∗ ∗ p* = 1×10*−*3, and QW vs. NREM *∗ ∗ p* = 9×10*−*3. **C)** Comparison of Hyperedge Centrality across different iEEG frequency bands. The Kruskal-Wallis test results indicate significant differences in the Delta band: Wake vs. NREM *∗ ∗ ∗ ∗ p* = 9×10*−*4 and REM vs. NREM *∗ ∗ ∗ ∗ p* = 3×10*−*5; and in the Theta band: AW vs. NREM *∗ p* = 3×10*−*2, based on Dunn’s multiple comparisons test. **D** Vertex Degree Centrality across different states. It represents the connections between two nodes that are linked across three or more frequency bands. **D)** Distance measures used in this study: Spectral Entropy, Hyperedge Centrality, and Vertex Centrality (from left to right).

For each sleep state, 100 epochs were selected from the total recordings. For each epoch, the signals were filtered into the following frequency bands: *δ* [1, 4] Hz, *θ* [4, 8] Hz, *α* [8, 12] Hz, *σ* [12, 15] Hz, *β* [15, 24] Hz, and *γ* [38, 42] Hz. The hypergraph construction method described in Section 3 was then applied. Finally, the quantifiers detailed in Section 2.2 were calculated for each hypergraph, sleep state, and rat. The average values were calculated across the epochs, resulting in a set of nine values (one per animal) for each sleep state.

Figure 2B shows that spectral entropy is highest during wake and REM sleep, both states being clearly distinct from QW and NREM. Interestingly, there were no significant differences between wake and REM, while the QW state displayed intermediate entropy, and NREM showed the lowest.

For the Hyperedge centrality analysis (Fig. 2C) in the delta band revealed significantly higher connectivity in wake and REM compared to NREM, though no differences were found between wakefulness, REM, and QW in other bands. The theta band, however, showed higher wakefulness values compared to NREM.

To study vertex degree centrality, we display the vertex that are connected in more than 3 bands (Fig. 2D). We can see visual and olfactory regions showed strong connectivity during both wakefulness and REM. In contrast, NREM displayed a marked reduction in connectivity, particularly the loss of connections in the visual areas (V2l), while olfactory bulb (OB) connectivity with (S1r) intensified. These findings align with prior studies, highlighting the role of these regions in maintaining cognitive function during wake and REM.

Figure 2E reveals clear distinctions between sleep states in terms of spectral, hyperedge, and vertex centrality distances. The wake and REM states are closely aligned, with the largest distance observed between NREM and wake/REM. These metrics effectively capture the differences in neural dynamics across the sleep cycle.

### 4.2 Differentiating Patients with Various Types of Epilepsy

The second example, where our method was applied, involved a study and comparison of brain dynamics in patients with different types of epilepsy, including those between interictal events. Understanding the dynamics of various seizure types is essential for improving diagnosis, treatment, and seizure detection technologies. It enables personalized therapies, prevents misclassification, and reveals the underlying neural mechanisms of seizures. Ultimately, this contributes to better management of epilepsy and advancements in research and treatment.

The dataset used in this analysis is an open-source electroencephalogram (EEG) dataset from [25]. It contains recordings from patients at the Epilepsy Monitoring Unit of the American University of Beirut Medical Center. These patients were diagnosed with focal epilepsy and underwent pre-surgical evaluations with long-term video-EEG monitoring to assess their suitability for epilepsy surgery. During this evaluation, antiepileptic medications were discontinued to capture natural seizures. EEG data were recorded using 19 scalp electrodes, following the 10 − 20 electrode placement system, with a sampling rate of 500 Hz. The datasets were pre-processed with band-pass filtering between 1/1.6 Hz and 70 Hz, excluding the 50 Hz power line frequency. Some channels were excluded due to artifacts.

The signals were segmented into one-second intervals, resulting in a 19 × 500 matrix per segment. Expert physicians classified the signals into four categories: Interictal Data (INIC): Resting-state signals without seizures. Complex Partial Seizures (CPS): Seizures with clear focal ictal EEG activity and clinical symptoms, such as altered awareness or automatisms. Electrographic Seizures (ES): Seizures detected only by EEG, with no observable clinical symptoms, but clear seizure activity on the EEG. Video-Detected Seizures with No Visual Change on EEG (VSNV): Clinically visible symptoms captured on video, but without corresponding EEG changes, suggesting non-epileptic activity or activity is not detectable by surface EEG. For each segment, the signals were filtered into the following frequency bands: *δ* [1, 4] Hz, *θ* [4, 8] Hz, *α* [8, 12] Hz, *σ* [12, 15] Hz, *β* [15, 24] Hz, and *γ* [38, 42] Hz. To each segment, we applied the hypergraph construction method described in Section 3 and, using the quantifiers from Section 2.2.

Figure 3B shows the entropy analysis across the different seizure types. Both the CPS and ES groups exhibited significantly higher entropy than the INIC and VSNV groups. However, no significant difference was found between CPS and ES or between VSNV and INIC.

**Figure 3:**
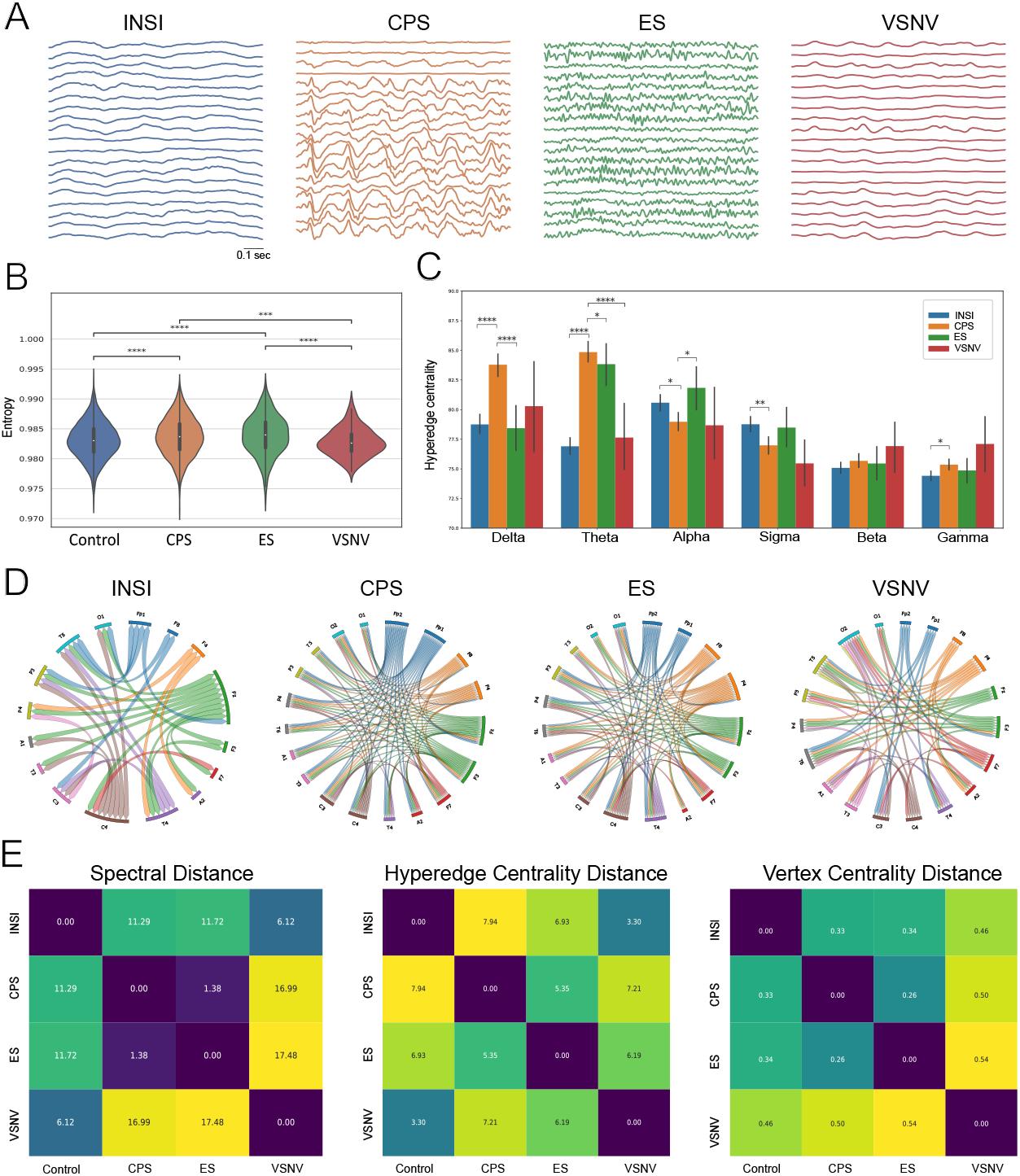
A study of hypergraph analysis in patients with different types of epilepsy: (INIC) resting-state signals without seizures, (CPS) complex partial seizures, (ES) electrographic seizures, and (VSNV) video-detected seizures with no visual change on EEG. **A)** EEG raw data recording example corresponding of each group. **B)** Spectral entropy analysis between different states. The violin’s plot shows the average entropy values over *N*_*INIC*_ = 3895, *N*_*CPS*_ = 3034, *N*_*ES*_ = 705, and *N*_*V SNV*_ = 111 epochs (Statistical test between states was made by Kruskal-Wallis test, for INIC vs CPS *∗ ∗ ∗ ∗ p* = 1.0×10^−7^, INIC vs ES *∗ p* = 3.0×10^−6^; CPS vs VSNV *∗ p* = 2.5×10^−2^, ES vs VSNV *∗ ∗ ∗ p* = 2.0×10^−4^ in Dunn’s multiple comparisons test). **C)** Comparison between Hyperedges Centrality for the different EEG bands. For Delta band (Kruskal-Wallis test, for Delta band INIC vs CPS *∗ ∗ ∗ ∗ p* = 1.6×10^−12^ and CPS vs ES *∗ ∗ ∗ ∗ p* = 5.0×10^−6^, for Theta Band INIC vs CPS *∗ ∗ ∗ ∗ p <* 1.0×10^−5^, CPS vs ES *∗ ∗ ∗ ∗ p* = 2.0×10^−6^ and CPS vs VSNV *∗ p* = 3.0×10^−2^, Alpha band INIC vs CPS *∗ p* = 3.0×10^−2^ and CPS vs ES *∗ p* = 2.5×10^−2^, Sigma band INIC vs CPS *p* = *∗ ∗* 1.3×10^−3^, and Gamma band INIC vs CPS *∗ ∗ p* = 1.7×10^−2^ in Dunn’s multiple comparisons test). **D)** Vertex degree centrality analysis, plotting the relationship between channels that, on average, exhibit connectivity across equal to or more than 3 frequency bands. **E)** Distance measures used in this work: Spectral Entropy, Hyperedge Centrality, and Vertex Centrality, from left to right.

In order to study hyperedge centrality we can see in the Delta band (Fig. 3C), the CPS group had the highest hyperedge centrality values, with a statistically significant difference compared to the INIC and ES groups. A trend toward higher values was observed in the VSNV group, though this was not statistically significant. Regarding theta band, the CPS group exhibited significantly higher values than the INIC, ES, and VSNV groups. In the Alpha band, the INIC group had higher values than the CPS group, while the ES group had significantly higher values than the CPS group. In the Sigma band, significant differences were only observed between the INIC and CPS groups. No significant differences were noted in the Beta band. Finally, in the Gamma band, the CPS group showed higher values compared to the INIC group.

For the Vertex degree centrality analysis, the same criteria as those employed in the previous example were applied. The circular plots illustrate the interactions between channels that are connected in three or more bands (Fig. 3C). In the INIC group, three electrodes (Fp2, T6, O2) did not meet the criterion of connecting across an average of more than 3 frequency bands. The Fz and C4 electrodes exhibited the highest number of connections with other electrodes in this group. In contrast, all three epileptic groups (CPS, ES, VSNV) displaying more connections between channels than in the INIC group. The CPS group demonstrated broad connectivity across all electrodes, with particular strength in the frontal regions. The ES group showed a similar pattern, but had fewer connections between electrodes Fp2 and Fp1 compared to other regions. The VSNV group had fewer connections, especially in electrodes Fp2, Fp1, and Fz, compared to the other epileptic groups.

Figure 3E illustrates the three distances employed in this study across the three seizure groups and the interictal events. The spectral distance analysis showed that the CPS and ES groups were the closest in terms of spectral similarity, while both were significantly different from the VSNV and INIC groups. The VSNV and INIC groups exhibited smaller distances between them. The hyperedge centrality distance showed that VSNV and INIC were the most similar groups, unlike the spectral distance measure, which was able to distinguish between CPS and ES. Lastly, vertex centrality mirrored the spectral distance metric, though the relative distance values between groups were smaller.

This analysis highlights distinct patterns of brain dynamics across different seizure types, emphasizing the potential of hypergraph-based methods in revealing underlying neural mechanisms and offering new insights for the diagnosis and treatment of epilepsy.

### 4.3 Dynamic Seizure Detection

The third case study demonstrates the application of our method to the analysis of multidimensional electrophysiological signals. The objective was to detect dynamic changes occurring during the onset and cessation of seizures in two patients diagnosed with generalized epilepsy.

Two patients were analyzed: Patient 1, diagnosed with primary generalized epilepsy, and Patient 2, with secondary generalized epilepsy. Both patients underwent magnetoencephalography (MEG) recordings during inter-ictal (resting) and ictal (seizure) periods. Recordings were taken with a 144-channel MEG at a sampling rate of 625 Hz. An example of a single MEG channel recording is shown in Fig. 4A.

**Figure 4:**
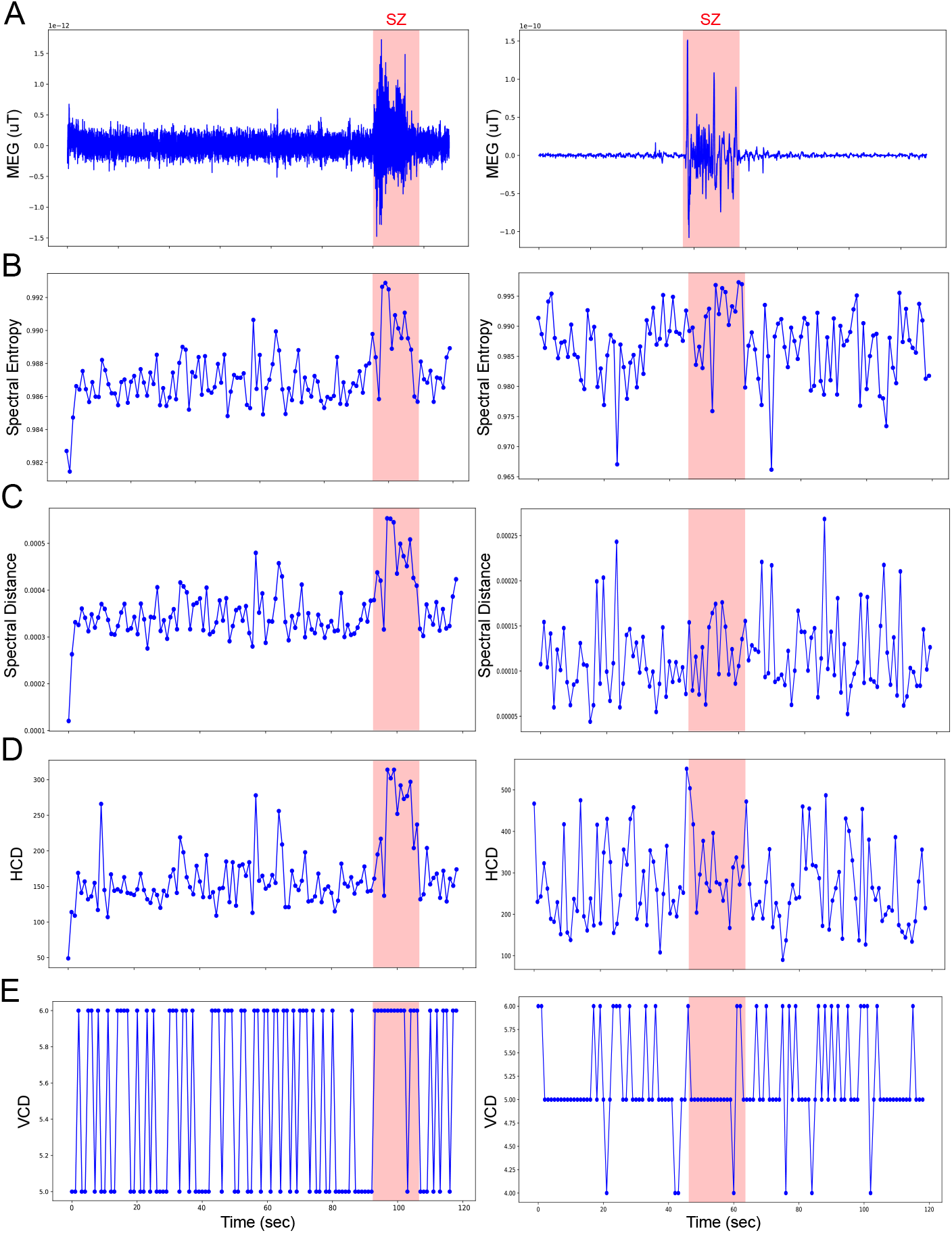
Study of changes in neural dynamics during two epileptic seizures. The left column corresponds to Patient 1, who suffers from generalized epilepsy (seizure type 1), and the right column corresponds to a patient with generalized epilepsy (seizure type 2). Each dot in the analysis represents a 1-second window of recorded data. The Shadow square defines the beginning and end of the seizure. **A)** Raw signal from a single channel of MEG data, where the red bar indicates the ictal period. **B)** Spectral entropy calculated over 120 windows. **C)** Spectral distance between the first window and subsequent windows. **D)** Similar to C, but using hyperedges centrality distance (HCD). **E)** Similar to C, but using vertex centrality distance (VCD).

Our primary aim was to assess changes in our quantifiers, specifically entropy, during the inter-ictal, ictal and postictal period. To do this, we employed multilayer hypergraph theory and its corresponding quantifiers to capture the dynamics of the MEG signals. We reduced the dimensionality of the data by selecting 42 of the 144 MEG channels for each Patient 2, ensuring a uniform distribution across the cerebral cortex. The original MEG signal was divided into 120 segments, each 1 second in length (625 data points per segment). For each segment, a multilayer hypergraph was constructed, as described in the Section 3, resulting in 120 hypergraphs for the entire recording duration. The frequency bands used for signal filtering and layer construction were *δ*[1, 4] Hz, *θ*[4, 8] Hz, *α*[8, 12] Hz, *β*_*low*_[12, 16] Hz, *β*_*high*_[20, 24] Hz, and *γ*[38, 42] Hz.

Figure 4B presents the results for entropy across all 120 windows for both patients. For Patient 1, during the basal state (0–95 seconds), the entropy values fluctuated within the range of [0.984, 0.988]. At approximately 97 seconds, corresponding to seizure onset, a sharp and abrupt increase in entropy was observed. The values then stabilized at a higher range between 0.990 and 0.992 during the seizure. After the seizure ended at 110 seconds, the entropy values decreased, resembling the pre-seizure baseline levels. In Patient 2, the baseline period (0–43 seconds) showed more variability in entropy, with values oscillating between [0.97, 0.99]. At 44 seconds, with the onset of the seizure, entropy increased, starting around 0.99 and reaching 0.995 by the end of the seizure at 65 seconds. Interestingly, there was a noticeable decrease in entropy values before the seizure, and after the seizure, the postictal entropy displayed variability similar to the inter-ictal values. These findings suggest a marked change in brain activity during the seizure, highlighting entropy as a potential biomarker for detecting seizure onset and recovery.

In Figures 4C, D, and E, we further analyze the data by calculating distances between hypergraphs in different time windows. For this analysis, we fixed the hypergraph from the first window (*H*_1_) and computed the distance between *H*_1_ and the hypergraphs from the subsequent windows (*H*_2_, *H*_3_, …, *H*_120_).

Figure 4C, shows the spectral distance, Patient 1 exhibits minimal distance between windows during the baseline period. However, at seizure onset, a significant increase in spectral distance occurs, which persists until the end of the seizure. Afterward, during the postictal period, the distance returns to baseline levels. In contrast, Patient 2 shows no significant changes in spectral distance throughout the recording.

Figure 4D shows that the hypergraph distance behaves similarly to the spectral distance for both patients. For patient 1, the hypedges distance increases during the epileptic seizure and returns to baseline levels once the seizure ends. Patient 2, however, exhibits a spike at the onset of the seizure, but the values quickly return to the interictal state. Figure 4E, depicts vertex centrality distance where different pattern emerges. Non-significant changes can be viewed in patient 2.

## 5 Discussion

Higher-Order Interaction (HOI) theory, as applied in this study, provides a powerful framework for capturing the complex, multi-dimensional relationships inherent in neuroscientific data. Unlike traditional graph-based methods that rely on pairwise interactions, HOI enables the analysis of intricate, non-linear dependencies across multiple components. This approach proved particularly effective in addressing the high-dimensional challenges posed by modern neurophysiological datasets, such as EEG and MEG signals [1]. A commonly used method for analyzing higher-order interactions (HOIs) in different systems is hypergraph theory. This approach offers a versatile and powerful framework for interpreting complex, multi-dimensional relationships across a variety of fields. Here, we applied hypergraph theory to analyze HOI in multidimensional electrophysiological signals. The hypergraphs were built from simple graphs generated through connectivity analysis across different electrophysiological frequency bands.

The analysis of sleep stages using hypergraph theory reveals valuable insights into brain dynamics. By examining spectral entropy, hyperedge centrality, and vertex centrality, we can distinguish between different sleep states. Wakefulness and REM sleep exhibit similarities in entropy and vertex centrality, a finding consistent with previous studies linking these states to higher cognitive processing and elevated brain activity [26, 27]. Interestingly, wakefulness and REM both show higher entropy values compared to NREM sleep, especially in the deeper stages like slow-wave sleep (SWS), which is associated with more synchronized, low-frequency brain waves such as delta waves [28].This reduced neural complexity is associated with the restorative processes, energy conservation and synaptic pruning of NREM sleep [29, 30].

Further analysis of vertex degree centrality reveals distinct patterns of connectivity across brain regions during different sleep stages. Visual and olfactory regions, for example, play prominent roles in wakefulness and REM, consistent with previous findings [31, 32, 33]. In contrast, NREM displays reduced connectivity in these regions, consistent with earlier studies indicating simplified neural communication during these restorative phases [34, 35]. Next, the distance between hypergraphs offers an additional layer of insight. The analysis of spectral, hyperedge, and vertex centrality distances across different sleep states reveals key differences in neural dynamics. For instance, the close similarity between wakefulness and REM reflects shared neural activity patterns, consistent with prior research [36]. In contrast, the larger distance between NREM and both wakefulness and REM suggests pronounced differences in neural function during NREM, as has been shown in earlier studies [37]. An interesting observation is that in rats, even experienced electrophysiologists sometimes find it challenging to distinguish quiet wakefulness (QW) from a light sleep state. In other words, some epochs might represent transitional states. However, in this study, we have demonstrated that QW can be reliably differentiated from both REM and NREM sleep across all quantifiers.

In our second example, we analyzed different types of seizures. Clinically, the differentiation between seizure types—Complex Partial Seizures (CPS), Electrographic Seizures (ES), and Video-Detected Seizures with No Visual Change on EEG (VSNV)—is well-established [38, 39, 40]. However, comparative studies exploring the dynamics and collective behavior of signals in these categories remain limited, hindering the contextualization of our results. Our analysis found that entropy and hyperedge centrality metrics provide important insights into the neural mechanisms underlying these seizure types. Spectral entropy, for example, was significantly higher in the CPS and ES groups compared to the INIC (interictal) and VSNV groups. Higher entropy in the Laplacian indicates a more uniform distribution of eigenvalues, suggesting more homogeneous connectivity between frequency bands. This aligns with previous research showing reduced variability in neural dynamics during seizures [41]. The analysis of hypergraph distances further enhanced the discrimination between seizure groups. Spectral distance, for instance, effectively distinguished between CPS and ES, while hyperedge centrality highlighted similarities between VSNV and INIC groups. These complementary measures provide a more comprehensive understanding of seizure dynamics, potentially improving diagnostic accuracy.

In the context of dynamic seizure detection, the results demonstrate that multilayer hypergraph analysis can effectively characterize the dynamic changes in MEG signals. The use of entropy and the three distances introduced provided valuable insights into the temporal evolution of brain activity during the different phases of the seizure: basal (resting), ictal (seizure), and postictal (recovery) periods. In both patients, entropy increased significantly during seizure onset, consistent with prior research suggesting that seizure activity often results in higher and more uniform connectivity across frequency bands. This is reflected in a more normal distribution of the eigenvalues of the Laplacian. In Patient 1, the abrupt rise in entropy during the ictal phase, followed by a return to near-baseline levels post-seizure, indicates a clear shift in cortical dynamics during the seizure event. This pattern suggests that entropy may serve as a reliable indicator for detecting seizure onset and cessation in primary generalized epilepsy. In Patient 2, the increase in entropy was less pronounced but still observable, suggesting that entropy can also capture dynamic changes in secondary generalized epilepsy. The analysis of hypergraph distances further supports these findings. The three distance measures effectively captured changes in network dynamics for Patient 1. The persistence of elevated distances throughout the seizure and their return to baseline postictally reinforces the idea that seizures induce large-scale increases in the brain’s functional connectivity [42]. For Patient 2, however, these distance measures did not detect significant changes during the seizure event. This discrepancy may be explained by the differing nature of seizures in primary and secondary generalized epilepsy. In primary generalized epilepsy, seizures originate simultaneously in both hemispheres of the brain, often due to genetic factors, and are characterized by a sudden loss of consciousness [43]. In contrast, secondary generalized epilepsy begins with a focal seizure, which then spreads to involve both hemispheres, typically resulting from structural brain abnormalities, and is marked by more gradual, evolving seizure patterns [43]. These differences in the nature of epilepsy may explain the variations in the behavior of quantifiers between the two patients.

In this study, we employed a single connectivity metric between bands, Phase Locking Index (PLI), as it is optimally suited for signals that have been filtered into frequency bands. Nevertheless, alternative connectivity metrics, including coherence, mutual information, and cross-entropy, could be employed for each frequency band. Furthermore, it is thought that this methodology can be extended to the analysis of fMRI signals, despite the fact that fMRI signals cannot be separated into frequency bands in the same way as EEG or MEG. Alternatively, distinct layers may be generated through the use of different correlation metrics between regions of interest (ROIs), including Pearson correlation, mutual information, or some divergence. Each metric offers a distinct perspective on the interconnections between voxels. Subsequently, hypergraphs can be constructed and analysed in a manner analogous to that employed in this study.These ideas will be implemented in future work. Also, in the future, we intend to develop and apply additional metrics to hypergraphs in order to ascertain whether further information can be extracted. In particular, our objective is to examine the Laplacian spectrum and associated metrics that may offer more precise insights than entropy, with a focus on local variations in Laplacian eigenvalues. Furthermore, we intend to apply these metrics to other challenges, such as the early detection of Alzheimer’s or Parkinson’s disease through the analysis of EEG or MEG signals. The hypergraph method introduced in this work allows for the extraction of higher-order information, which could be used as quantifiers in deep learning models. These are areas we plan to explore in the near future.

## 6. Conclusion

This study demonstrates the value of Higher-Order Interaction (HOI) theory, particularly through hypergraph analysis, in uncovering complex, multidimensional patterns within neuroscientific data. By applying hypergraph theory to analyze brain dynamics across different states, such as sleep and seizure phases, the study reveals meaningful distinctions in neural connectivity and entropy that traditional pairwise methods might overlook. This insight highlights HOI’s capacity to refine our understanding of neural behavior, offering a robust framework for future studies in consciousness and potentially improving diagnostic precision in neurological conditions.

## Acknowledgements

This publication was supported by grand # MinCyT-FonCyT PICT-2019 N° 01750 PMO BID; grant CONICET-PUE-IMAL # 229 201801 00041 CO; grant CONICET-PIP-2021-2023-GI #11220200101940CO and grant UNL-CAI+D # 50620190100070LI

## Notes

### Competing Interest Statement

The authors have declared no competing interest.

